# Transcriptional profiling reveals conserved and species-specific plant defense responses during the interaction of the early divergent plant *Physcomitrium patens* with *Botrytis cinerea*

**DOI:** 10.1101/2020.10.29.361329

**Authors:** Guillermo Reboledo, Astrid Agorio, Lucía Vignale, Ramón Alberto Batista-García, Inés Ponce De León

**Affiliations:** Departamento de Biología Molecular, Instituto de Investigaciones Biológicas Clemente Estable, Montevideo, Uruguay; Centro de Investigación en Dinámica Celular. Universidad Autónoma del Estado de Morelos, Cuernavaca, México

**Author notes:** corresponding autor.

**Keywords:** *Physcomitrium patens*, *Botrytis cinerea*, transcriptome, defense genes, orphan genes

## Abstract

Bryophytes were among the first plants that colonized earth and they evolved key defense mechanisms to counteract microbial pathogens present in the new environment. Although great advances have been made on pathogen perception and subsequent defense activation in angiosperms, limited information is available in early divergent land plants. In this study, a transcriptomic approach uncovered the molecular mechanisms underlying the defense response of the bryophyte *Physcomitrium patens* against the important plant pathogen *Botrytis cinerea*. A total of 3.072 differentially expressed genes were significantly affected during *B. cinerea* infection, including genes encoding proteins with known function in angiosperm immunity and involved in pathogen perception, signaling, transcription, hormonal signaling, metabolic pathways such as shikimate and phenylpropanoid, and proteins with diverse role in defense against biotic stress. Similarly as in other plants, *B. cinerea* infection leads to downregulation of genes involved in photosynthesis and cell cycle progression. These results highlight the existence of evolutionary conserved defense responses to pathogens throughout the green plant lineage, suggesting that they were probably present in the common ancestors of land plants. Moreover, several genes acquired by horizontal transfer from prokaryotes and fungi, and a high number of *P. patens*-specific orphan genes were differentially expressed during *B. cinerea* infection, indicating that they are part of the moss immune response and probably played an ancestral role related to effective adaptation mechanisms to cope with pathogen invasion during the conquest of land.

**Key Message:** Evolutionary conserved defense mechanisms present in extant bryophytes and angiosperms, as well as moss-specific defenses are part of the immune response of the early divergent land plant *Physcomitrium patens*.

## Introduction

Bryophytes, including mosses, liverworts, and hornworts, are small non-vascular plants that were among the first plants that colonized land approximately 450 million years ago (Mya). Due to the transition from an aquatic to a terrestrial environment, bryophytes acquired adaptation mechanisms to cope with different kinds of abiotic stresses, including desiccation stress, variations in temperature, and UV-B radiation, as well as defense mechanisms against microorganisms present in the air and soil (Rensing et al. 2008; Ponce de León and Montesano 2017). Bryophytes diverged before vascular plants appeared and represent therefore excellent models to reveal ancient defense mechanisms against pathogens. However, in contrast to the vast information available in angiosperms-pathogen interactions, only few evidences show the recognition of pathogens and subsequent activation of defense responses in early land plants such as mosses and liverworts.

Plants are in permanent contact with microbial pathogens, including fungi, bacteria, viruses and oomycetes. In order to detect the presence of the pathogen, angiosperms have evolved complex signaling and perception pathways. Microorganisms have carbohydrate- and protein-based signals that are essential for microbial survival, for example flagellin or chitin, classified as pathogen-associated molecular patterns (PAMPs) (Boller and Felix 2009). Based on their conservation, and the fact that they are not produced in plant cells, plants synthesize different plasma membrane localized pattern recognition receptors (PRRs), including receptor-like kinases (RLKs), that recognize PAMPs to control plant immunity. In response to PAMPs, plants trigger a defense response called PAMP-triggered immunity (PTI) or basal resistance, which is the first level of defense that restricts pathogen infection in most plant species (Jones and Dangl 2006). Pathogens adapted to their host plants have evolved strategies to interfere and inhibit plant defense by the action of pathogen-secreted virulence factors known as effectors that target key PTI components (Boller and Felix 2009). Plants have a second layer of immune system consisting of receptors encoded by resistance (R) genes to detect directly or indirectly the effector proteins leading to effector-triggered immunity (ETI) (Jones and Dangl 2006). Both PTI and ETI result in a rapid burst of extracellular reactive oxygen species (ROS), activation of mitogen-activated protein kinases (MPKs), increase in hormone synthesis, and induction of genes with different roles in plant defense (Jones and Dangl 2006). ETI is often accompanied by a hypersensitive response (HR), a type of programmed cell death that restricts the pathogen to the site of infection.

In bryophytes, most of the knowledge related to defense activation has been generated during the interaction of microbial pathogens with the moss *Physcomitrella patens* (Ponce de León and Montesano 2017), which has been recently renamed as *Physcomitrium patens* (*P. patens*) (Rensing et al. 2020). Pathogenic bacteria, fungi and oomycetes infect *P. patens* tissues leading to maceration, necrosis and cell death (Ponce de León 2011). *P. patens* senses the presence of fungal pathogens by perceiving chitin through the receptor CERK1 and consequently activates a MPKs signaling cascade, which is necessary for plant immunity (Bressendorff et al. 2015). Hormonal levels increases after pathogen colonization in moss tissues, including salicylic acid (SA), abscisic acid (ABA), auxin and the precursor of jasmonic acid (JA), cis-oxophytodienoic acid (OPDA), since JA is not synthesized in bryophytes (Ponce de León et al. 2012; Mittag et al. 2015). Expression levels of several genes encoding proteins with different roles in defense are induced in *P. patens*-infected tissues, including pathogenesis-related proteins (PRs) with putative antimicrobial activities such as PR-1 and PR-10 (Ponce de León et al. 2007; Castro et al 2016). After pathogen colonization, *P. patens* activates cell wall reinforcement by incorporation of phenolic compounds, callose deposition and induction of genes encoding Dirigent (DIR) proteins involved in the synthesis of lignin-like compounds (Ponce de León et al. 2012; Reboledo et al. 2015; Alvarez et al. 2016). Recent studies in the liverwort *Marchantia polymorpha* (*M. polymorpha*) have shown that infection with the oomycete *Phytophthora palmivora* (*P. palmivora*) induces a defense response leading to increased expression of transcription factors (TFs), PRs and genes related to the phenylpropanoid pathway, indicating the several plant defense mechanisms are conserved between liverwort and angiosperms (Carella et al. 2019).

*Botrytis cinerea* (*B. cinerea*) is a necrotrophic fungus that infects over 200 different plant species including *P. patens*, and causes maceration and necrosis of the plant tissues (Ponce de León et al. 2007). Several virulence factors, including toxins and cell wall degrading enzymes such as endopolygalacturonases and xylanases are important to cause disease (van Kan 2006). In response to *B. cinerea*, *P. patens* induces ROS production, activates an HR-like response, reinforces the cell wall, increases SA and OPDA synthesis and induces the expression of several genes with known defense functions (Ponce de León et al. 2007, 2012). To get more insights into *P. patens*-pathogen interactions, and since large scale transcriptional studies during pathogen infection are not available for mosses, we performed RNAseq profiling of *P. patens* during *B. cinerea* infection. Our results demonstrate the existence of an evolutionary conserved immune system among the plant lineage, where genes involved in perception, signaling, transcription, secondary metabolism and genes with diverse roles in defense against biotic stress, participate in the response of this early land plant to *B. cinerea* infection. Moreover, species-specific orphan genes participate in defenses against this pathogen, suggesting an ancestral role probably related to adaptation mechanisms to cope with pathogen invasion during the conquest of land.

## Material and methods

### Plant material and pathogen inoculation

*P. patens* Gransden wild type colonies were cultivated on solid BCDAT medium (Ashton and Cove 1977) under standard long-day conditions (22°C, 16-h light/8-h dark regime under 60–80 μmol m^2^ s^-1^ white light) for 3 weeks before spray inoculation with a *B. cinerea* 2×10^5^ spores/mL suspension or water (mock). Three time points corresponding to spore germination (4 hours post inoculation; hpi), germ tubes elongation (8 hpi), and moss cells colonization (24 hpi) were analyzed. Three independent biological replicates of tissue were harvested at each time point and treatment for RNA extraction, immediately frozen in liquid nitrogen, and stored at −80°C.

### *B. cinerea* staining

*B. cinerea* tissues were stained with 0.1% solophenyl flavine 7GFE 500 according to Ponce de Leon et al. (2012). Fluorescence microscopy was performed with an Olympus BX61 microscope (Shinjuku-ku, Japan). Photographs were taken at 4, 8 and 24 hpi.

### RNA extraction, cDNA library preparation and sequencing

Frozen samples were pulverized with a mortar and pestle and total RNA was extracted using the RNeasy Plant Mini Kit (Qiagen, Germany), including a RNase-Free DNase treatment in column (Qiagen, Germany), according to manufacturer’s protocol. RNA quality control, library preparation, and sequencing were performed at Macrogen Inc. (Seoul, Korea). RNA integrity was checked before library preparation using an Agilent Technologies 2100 Bioanalyzer (Agilent Technologies). Libraries for each biological replicate were prepared for paired-end sequencing by TruSeq Stranded Total RNA LT Sample Prep Kit (Plant) with 1 μg input RNA, following the TruSeq Stranded Total RNA Sample Prep Guide, Part # 15031048 Rev. E. Sequencing was performed on Illumina platform (Illumina, California, USA) by Macrogen Inc. (Seoul, Korea) to generate paired-end 101 bp reads, obtaining 41 to 64 M reads per sample with Q20 >98.31 %.

### RNA-seq processing

RNA-seq processing steps were done throw Galaxy platform (https://usegalaxy.org/). Raw reads quality was revised by FastQC software ver. 0.11.2 (http://www.bioinformatics.babraham.ac.uk/projects/fastqc/) and then preprocessed for both quality and adapter trimmings using Trimmomatic Version 0.38.0 software (Bolger et al., 2014). Additionally to default options, the following parameters were set: adapter sequence TruSeq3 (paired-ended, for MiSeq and HiSeq), always keep both reads of PE True, SLIDINGWINDOW: 4:15 HEADCROP:12 MINLEN:50. Trimmed reads were mapped to reference genomes of *P. patens* using Hisat2 software (Kim et al., 2015). All reads were mapped first against *P. patens* organelle genomes and rRNA sequences (mitochondrial NC_007945.1; chloroplast NC_005087.1; ribosomal HM751653.1, X80986.1 and X98013.1). The remaining reads were mapped against *P. patens* nuclear genome v3 (Lang et al., 2018), using Ppatens_318_v3.fa as the reference genome file and Ppatens_318_v3.3.gene_exons.gff3 as a reference file for gene models (https://phytozome.jgi.doe.gov/pz/portal.html), and concordant mapped read pairs were retained. The BAM files with read mapped in proper pair were obtained with Samtools View software ver. 1.9 and then sorted by name with Samtools Sort software ver. 2.0.3 (Li et al., 2009), for further analysis.

### Differential expression analysis

The reads were counted by the FeatureCounts software ver. 1.6.4 (Liao et al., 2013). Additionally, for default options, the following parameters were set: Stranded (Reverse), Count fragments instead of reads −p, Allow read to contribute to multiple features True, Count multi-mapping reads/fragments -M and Reference sequence file *P. patens* v3.3. Differential expression analyses were performed using EdgeR software ver. 3.24.1 (Robinson et al., 2010; Liu et al., 2015) with p-value adjusted threshold 0.05, p-value adjusted method Benjamini and Hochberg (1995) and Minimum log2 Fold Change 2. Counts were normalized to counts per million (cpm) with TMM method and low expressed genes filtered for count values ≥5 in three or more samples. In this study, a false discovery rate (FDR) ≤0.05 was used to determine significant differentially expressed genes (DEGs) between *B. cinerea* inoculated plants and mock, and expression values were represented by log2 ratio. Heatmaps with DEG data were constructed in GraphPad Prism software ver. 8.0.2.

### Quantitative real-time PCR validation of RNA-Seq Data

The expression level of twelve selected genes was analyzed to validate RNA-Seq results via quantitative reverse transcription PCR (RT-qPCR). cDNA was generated from 1 μg of RNA using RevertAid Reverse transcriptase (Thermo Scientific) and oligo (dT) according to the manufacturer’s protocol. RT-qPCR was performed in an Applied Biosystems QuantStudio 3 thermocycler using the QuantiNova Probe SYBR Green PCR Kit (Qiagen, Germany); mix proportions and cycling parameters were used as described in manufacturer’s instructions. Relative expression of each gene was normalized to the quantity of constitutively expressed Ubi2 gene (Le Bail et al., 2013), using the 2^-ΔΔCt^ method (Livak and Schmittgen 2001). Gene expression of *P. patens* inoculated tissues was expressed relative to the corresponding water-treated samples at the indicated time points, with its expression level set to one. Each data point is the mean value of three biological replicates. Student’s t-test was performed to determine the significance for quantitative gene expression analysis using GraphPad Prism software ver. 8.0.2. P-values <0.01 were considered statistically significant. Primer pairs used for qPCR analyses are provided in **SupplementaryTable S1**, in all cases amplification efficiencies was greater than 95%.

### GO enrichment analysis

Gene ontology (GO) and functional annotations were assigned based on *P. patens* v3.3 gene models (https://phytozome.jgi.doe.gov/pz/portal.html) and Blast2GO 5.2.5 software (Götz et al., 2008), through Blast and Interpro searches of homologs and protein domains. Additionally, to default options, for every step, the following parameters were set for Blast: e-value ≤ 1.0E-3 and species viridiplantae. DEG functional enrichment analysis was performed using OmicsBox tools (https://www.biobam.com/omicsbox/). GO terms with a FDR ≤0.05 were considered for our analysis. To summarize the long list of GO terms, redundant GO terms were removed with REViGO (reduce and visualize gene ontology) (Supek et al. 2011).

*P. patens* differentially expressed orphan genes were identified by selecting genes with no ancestry gene, according to phytozome.org available data, and no Blast (tblastn) hits in viridiplantae (e-value >1.0E-3) (Wilson el al., 2005).

## Results

### Global gene expression analysis of *P. patens* plants during *B. cinerea* infection

In previous studies, we have shown that *B. cinerea* infects *P. patens* tissues and that fungal biomass starts to increase after 8 hours post-inoculation (hpi), reaching high levels at 24 hpi (Ponce de León et al. 2012). To have more insights into *P. patens* defense response to *B. cinerea* infection, RNAseq profiling was performed at three different stages of the colonization event; spore germination (4 hpi), germ tubes elongation (8 hpi), and hyphae proliferation and tissue colonization (24 hpi) (**Fig. 1**). At 8 hpi, some plant cells start to respond to *B. cinerea*, evidenced by cell wall modifications detected with solophenyl flavine, and initial penetration attempts of some cells by infectious hyphae were visible (**Fig. 1d-e**). At 24 hpi, hyphae colonized *P. patens* cells and fungal structures could be clearly distinguished within *P. patens* cells (**Fig. 1f**). In our working conditions, disease symptoms were visible at 24 hpi evidenced by darkening of the tissues and heavy maceration at 48 hpi (**Fig. 1g-j**), which are typical symptoms caused by this pathogen in angiosperms.

**Fig. 1:**
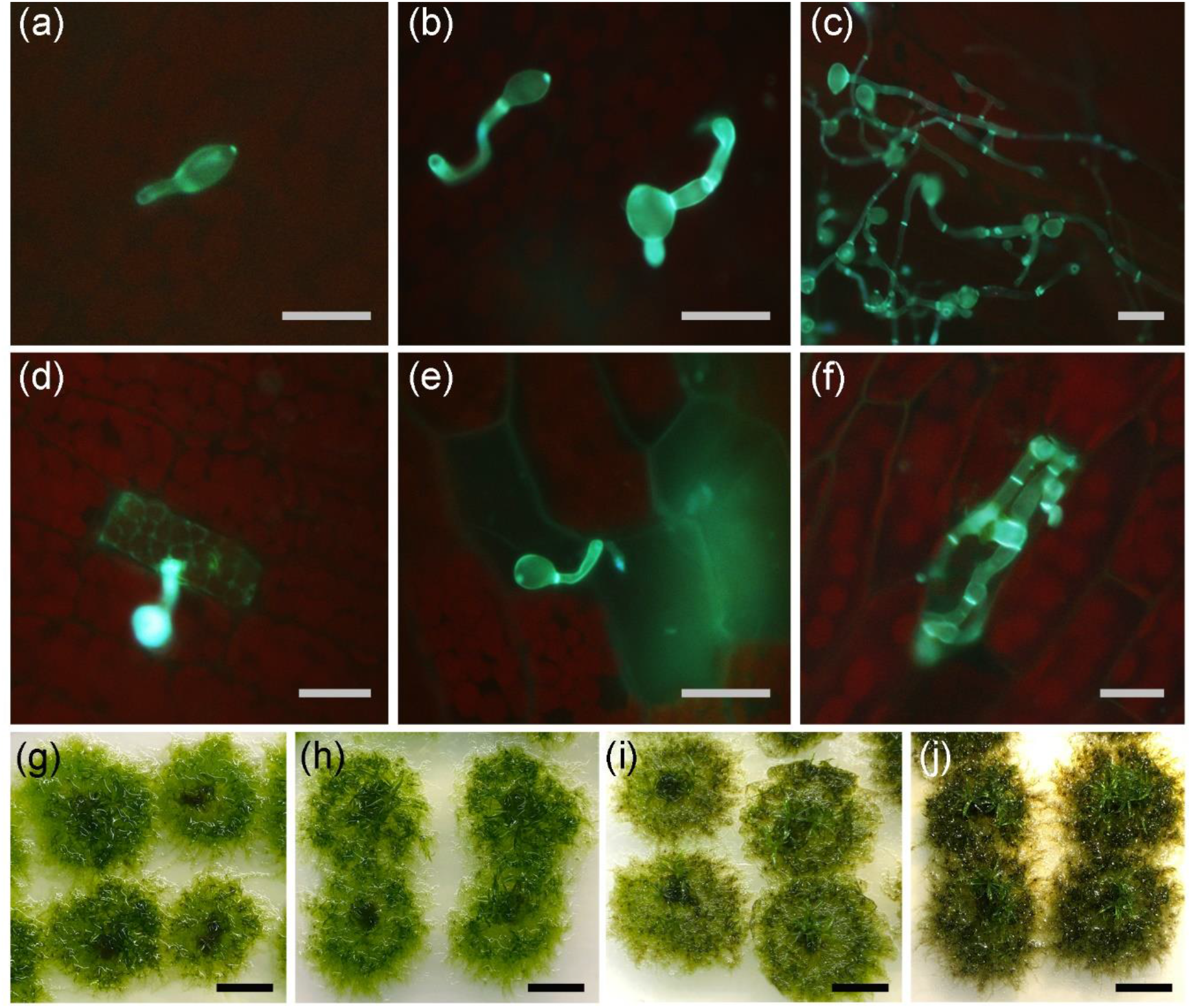
*B. cinerea* colonization and symptom development in *P. patens*. Infection progress during time is shown; **(a)** spore germination at 4 hpi, **(b)** Germ tubes elongation at 8 hpi, **(c)** proliferation of mycelium at 24 hpi. **(d)** Hyphal tip approaching a moss cell at 8 hpi, **(e)** initial penetration of infectious hypha, and **(f)** Hyphae growing within a moss cell at 24 hpi. Symptom development at 8 hpi **(h)**, 24 hpi **(i)**, 48 hpi **(j)**, and water-treated moss colonies **(g)** are shown. The scale bar in **a-f** represents 10 μm and 0,5 cm in **g-i**.

The transcriptomes of water-treated and *B. cinerea*-inoculated *P. patens* tissues were analyzed in three biological replicates. A total of 86.8-96.5% of the reads in the libraries mapped successfully to the genomes of *P. patens* (nuclear, chloroplast and mitochondria). Reads mapped uniquely to the *P. patens* nuclear genome (approximately 208 million reads) were used for further analyses (**Supplementary Table S2**). During the entire time course, 3.072 DEGs were identified, which represent 9,3 % of the 32.926 *P. patens* gene models (**Supplementary Table S3**). The three selected time points showed a clear distinguishable transcriptional profile in response to *B. cinerea* inoculation (**Fig. 2a**). Only 17 DEGs were identified at 4 hpi (all upregulated), while a massive transcriptional shift towards upregulation was observed at 8 hpi (1.032 upregulated versus 150 downregulated genes). At 24 hpi, transcriptional changes included a high number of upregulated genes (1.208) as well as downregulated genes (1.482). For this work, we defined the three time points as Early Response (ER; 4 hpi), Middle Response (MR; 8 hpi) and Late Response (LR; 24 hpi). Transcriptional dynamics show an overlap of DEGs in ER, MR and LR, and increased expression levels over time was evident. Of the 17 DEGs induced at ER, 16 and 9 were also upregulated at MR and LR, respectively (**Fig. 2b**). At MR and LR, 693 upregulated and 91 downregulated DEGs were in common, being the rest specific for each time point.

**Fig. 2:**
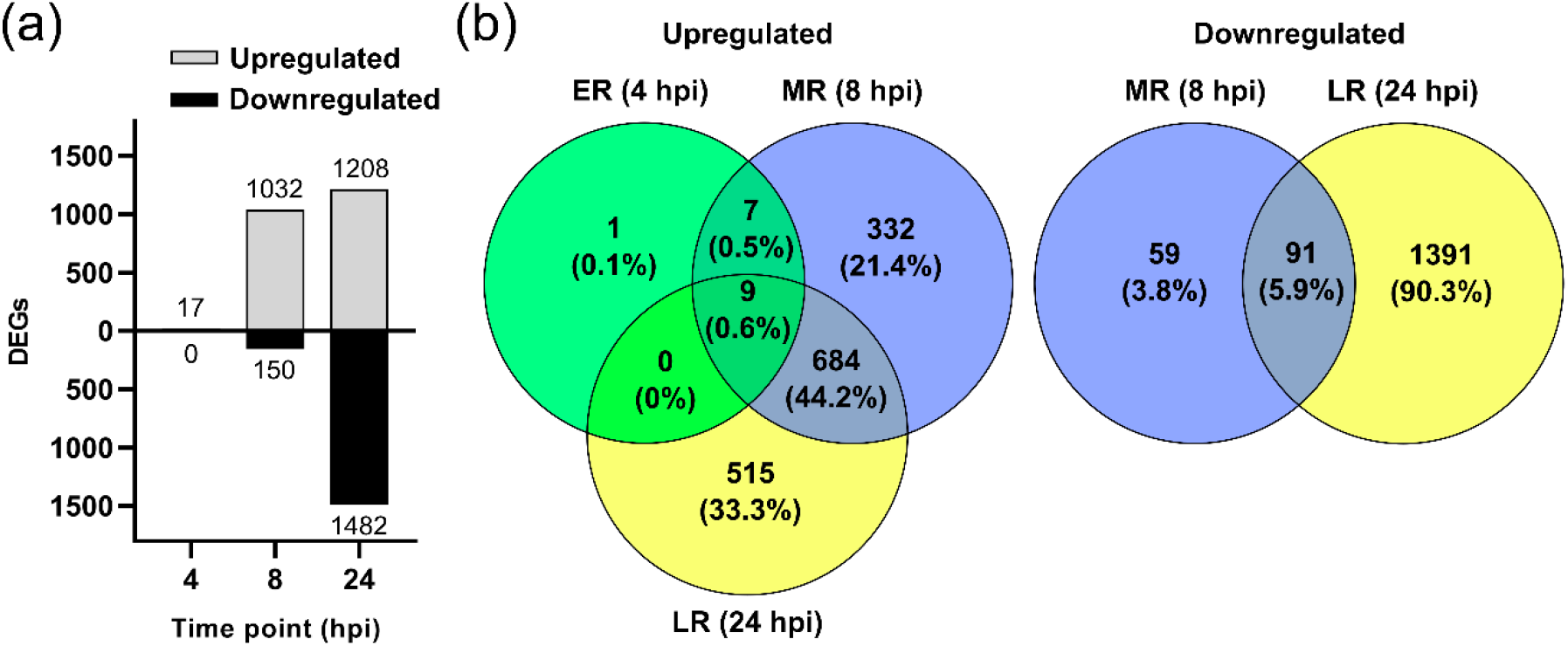
Differentially expressed *P. patens* genes during *B. cinerea* infection. **(a)** Number of differentially expressed genes (DEGs), up- and downregulated, in *P. patens* tissues inoculated with *B. cinerea* versus water-treated tissues at 4 hpi, 8 hpi and 24 hpi. (**b**) Venn diagram of *B. cinerea*-responsive *P. patens* genes at Early Response (ER; 4 hpi), Middle Response (MR; 8 hpi), and Late Response (LR; 24 hpi) showing overlap of expressed *P. patens* genes.

DEGs present at ER encoded proteins related to stress such as Late Embryogenesis Abundant proteins (LEAs), Hydrophobic protein RCI2, Low Temperature and Salt responsive protein (LTI), DNAJ protein, early light-induced protein (ELIP), transcription factor with an AP2 domain, peripheral-type benzodiazepine receptor (TSPO) and phenylalanine-ammonia lyase (PALs) (**Supplementary Table S4**). Genes encoding enzymes involved in signaling and synthesis of phosphatidic acid (PA) and sphingolipids, which are lipid messengers involved in plant response to biotic and abiotic stress, were also upregulated upon *B. cinerea* inoculation, including lipid phosphate phosphatase and glucosylceramidase. The high proportion of ER-inducible genes during MR and to a less extent during LR, accompanied by increased expression levels, indicate that they play a role in moss defense during initial stages of infection.

### Biological responses of *P. patens* to *B. cinerea* inoculation

Enrichment of Gene Ontology (GO) terms was performed to identify the biological processes mostly affected by *B. cinerea* infection (**Supplementary Table S5**). Many different GO terms were enriched underlining the wide extent of the plant response to this pathogen. Most of the top ten significantly enriched GO terms for the upregulated genes were similar at MR and LR, and included metabolic processes such as secondary metabolic process, phenylpropanoid, erythrose 4-phosphate/phosphoenolpyruvate family amino acid, L-phenylalanine and aromatic amino acid family metabolic process, and response to fungus, defense response and response to chemical (**Fig. 3**). Other upregulated defense-related GO terms present at MR and LR included response to abiotic stimulus, external stimulus, wounding, UV-B, and salt, as well as regulation of hormone levels and jasmonic acid biosynthetic process. Moreover, at LR, cell wall metabolic processes were enriched, including cell wall organization or biogenesis and regulation of cell wall macromolecule. Collectively, biotic, abiotic related processes and phenylpropanoids metabolism, accounted for more than 70% and 50% of the GO biological process (BP) terms identified in the up-regulated genes at MR and LR, respectively.

**Fig. 3:**
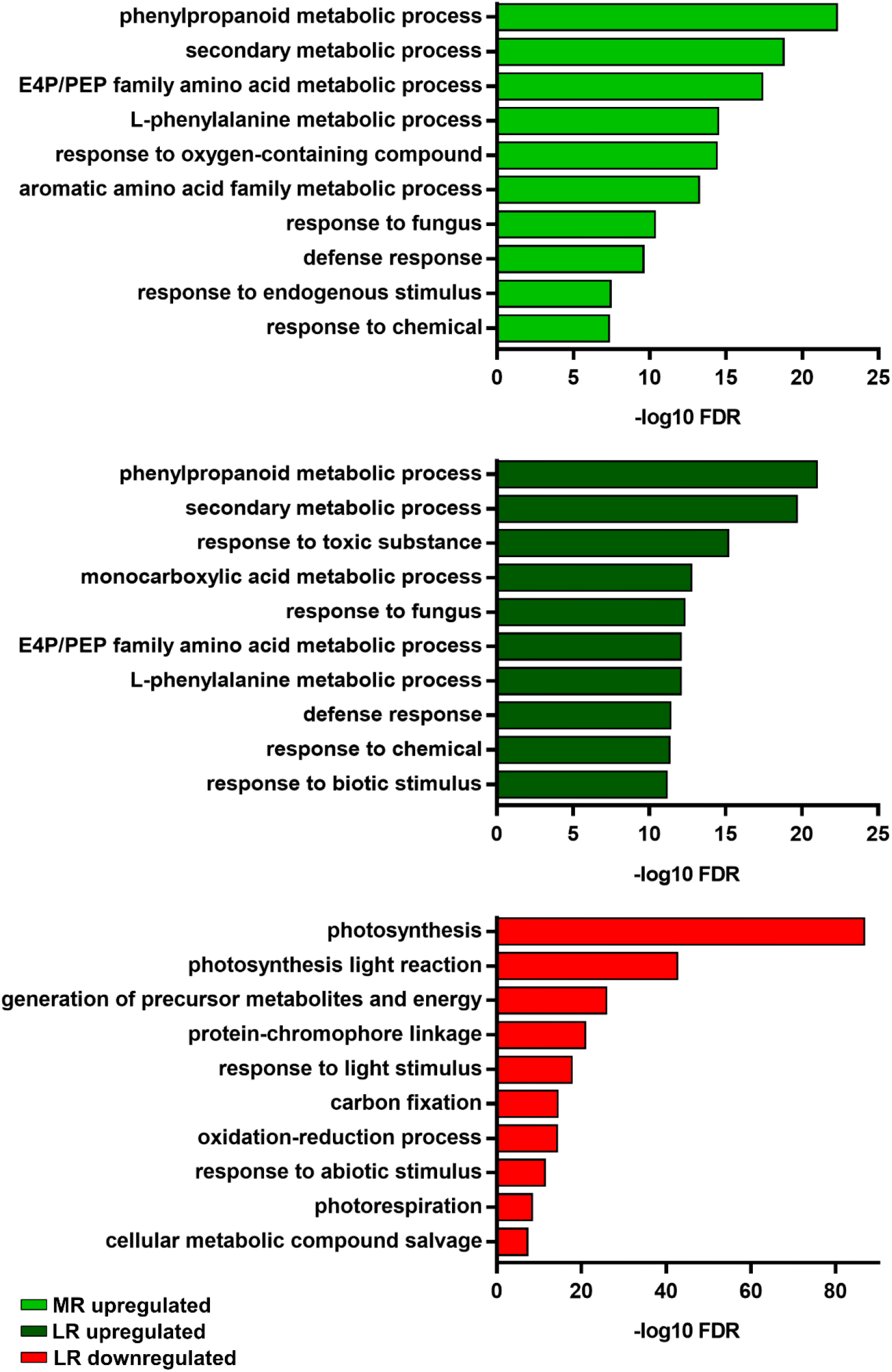
Enriched gene ontology (GO) biological process terms. The top 10 enrichment GO terms obtained by REViGO at MR and LR are shown and significance is expressed as -- log10 FDR. Abbreviation: E4P/PEP: erythrose 4-phosphate/phosphoenolpyruvate. See Supplementary Table S6 for complete information.

Significant enriched GO terms for downregulated genes were only identified at LR and were related to photosynthesis, including photosynthesis light reaction, generation of precursor metabolites and energy, protein-chromophore linkage, response to light stimulus, carbon fixation, photorespiration and chlorophyll metabolic process. Overrepresented downregulated DEGs included genes encoding chlorophyll binding proteins, proteins related to photosystem I and II, as well as many other proteins related to photosynthesis. Other significantly enriched GO terms were cell cycle process, cell cycle, nuclear division, cell division, chromosome segregation, and anaphase-promoting complex-dependent catabolic process. Within these overrepresented downregulated DEGs, genes encoded cyclins, cyclin-dependent kinases, centromeric proteins, cell division control proteins, kinesins, as well as other proteins involved in cell cycle progression and cell division (**Supplementary Table S5, S6**). Taken together, these results indicate that during MR, *P. patens* activates a general respond that encompasses several defense pathways that are maintained during LR, while at LR important processes such as photosynthesis, cell division and cell cycle progression are downregulated.

### Activation of the shikimate and phenylpropanoid pathways during *P. patens* defense against *B. cinerea*

GO term enrichment analysis (BP and Molecular Function; MF) and manual inspection of the identified *P. patens* DEGs showed a remarkable overrepresentation of genes involved in the shikimate and phenylpropanoid pathways (**Supplementary Table S5-S7**). Several of these genes were amongst the highest induced transcripts at MR and LR. In angiosperms, these pathways are responsible for the synthesis of secondary metabolites with diverse roles in defense, including reinforcement of the cell wall to halt pathogen progression, synthesis of SA and production of antimicrobial compounds (Dixon and Paiva 1995; Vogt 2010). In total, 12 and 13 genes of the shikimate pathway were upregulated at MR and LR, including gene members encoding eight different enzymes of the pathway (**Fig. 4a**; **Supplementary Table S7**). In addition, alteration of the phenylpropanoid pathway by *B. cinerea* starts at ER with increased expression of two *PALs* and continued during MR and LR with upregulation of 93 of 108 DEGs. DEGs encode PALs, cinnamic acid 4-hydroxylases (C4Hs) and 4-hydroxycinnamoyl-CoA ligases (4CLs), which play a crucial role in the formation of the substrate p-coumaroyl-CoA for both monolignol and flavanone biosynthesis. Other DEGs encode CHSs and chalcone isomerases (CHIs), as well as putative flavonol synthases (FLSs) and flavanone 3-hydroxylases (F3Hs), which are responsible for the production of flavonols in angiosperms. Several other DEGs encode enzymes involved in the biosynthesis of monolignols and lignin-like compounds such as hydroxycinnamoyl-CoA reductases (CCRs), caffeate O-methyltransferases (COMTs), and shikimate O-hydroxycinnamoyl transferases (HCTs). *B. cinerea* inoculation also led to significant increase in six DIR transcripts abundance, some of which were also upregulated *P. patens* tissues infected with other pathogens, and the encoded proteins are probably involved in coupling of monolignols to produce lignin-like compounds (Ponce de León and Montesano 2016). During *B. cinerea* infection, transcript levels of several genes encoding polyphenol oxidases (PPO) such as catechol oxidases and laccases, as well as peroxidases (Prx), increased significantly, which have a role in radical coupling of monolignols to form lignin and other phenolic compounds (Wang et al. 2013). Upregulation of *B*. cinerea-responsive genes encoding TFs TT2 (MYB), TT8 (bHLB), EGL1 and bHLH27 that are involved in flavonoid biosynthesis in other plants (Liu et al. 2015), were also observed. Gene expression values obtained from RNA-seq were validated using qPCR assay for 12 genes of these pathways. The results showed a strong correlation (MR; R2 =0.9648 and LR: R2=0.9465) between the results obtained with the two techniques (**Fig. 4b; Supplementary Table S8**). Since the complete phenylpropanoid pathway appeared for the first time in land plants and it likely evolved progressively by the recruitment of enzymes (Emiliani et al. 2009), our results suggest that phenylpropanoids played an important role in ancient plant defense mechanisms against pathogen infection.

**Fig. 4:**
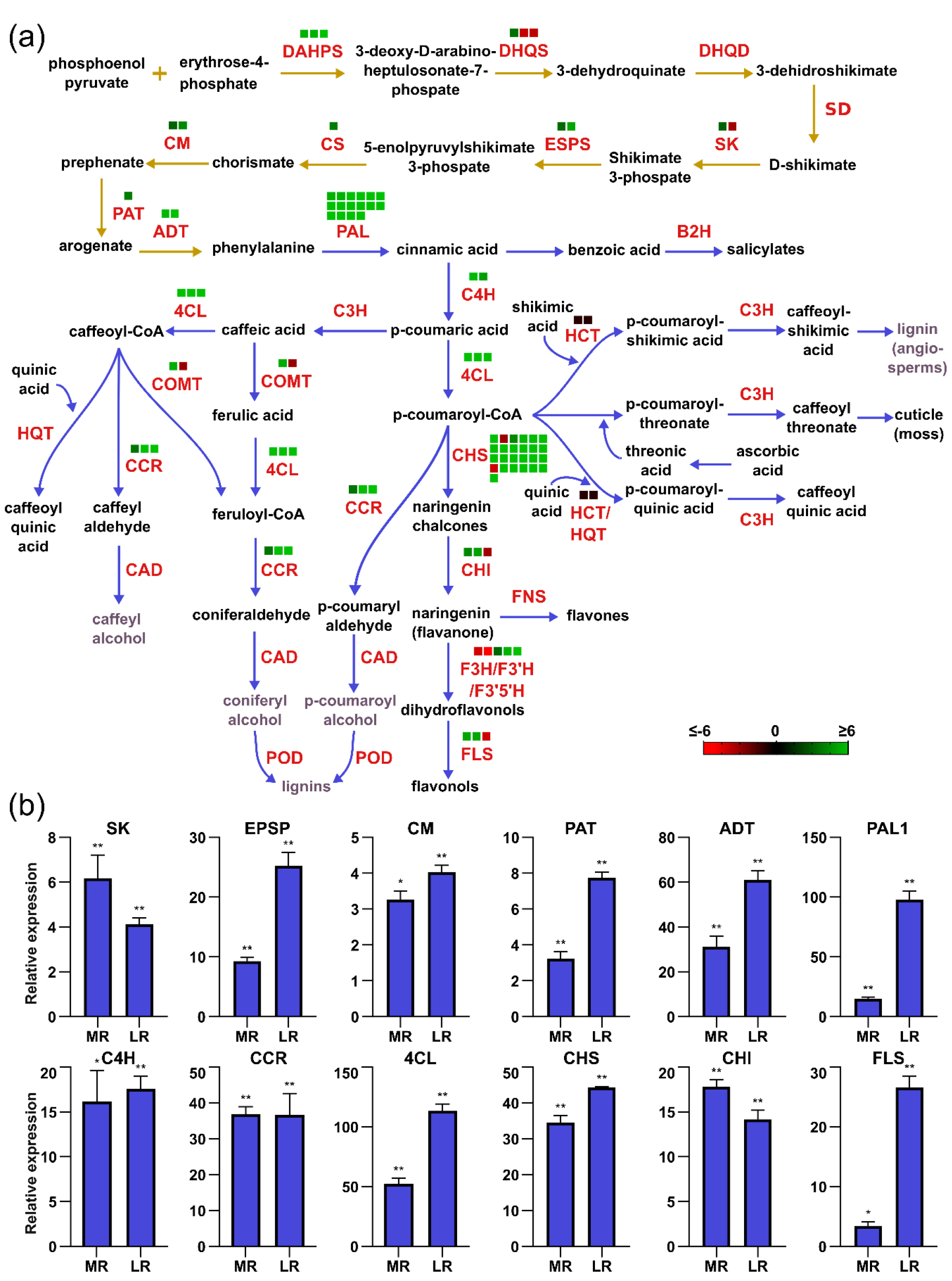
Activation of shikimate and phenylpropanoid pathways during *B. cinerea* infection. **(a)** Scheme of the shikimate and phenylpropanoid pathways. Heatmaps indicate log2 FC at LR. Abbreviations for the shikimate pathway: DAHPS, 3-deoxy-D-arabino-heptulosonate 7-phosphate synthase; DHQS, 3-dehydroquinate synthase; DHQD/SD, 3-dehydroquinate dehydratase/shikimate dehydrogenase; SK, shikimate kinase; ESPS, 3-phosphoshikimate 1-carboxyvinyltransferase; CS, chorismate synthase; CM, chorismate mutase; PAT, prephenate aminotransferase; ADT, arogenate dehydratase. Abbreviations for the phenylpropanoid pathway: PAL, phenylalanine ammonia-lyase; B2H, benzoic acid-2-hydroxylase; C4H, cinnamate-4-hydroxylase; 4CL, 4-coumarate CoA ligase; CHS, chalcone synthase; CHI, chalcone isomerase; F3H, flavanone 3-hydroxylase; F3’H, flavonoid 3’-hydroxylase; F3’5’H, flavonoid 3’,5’-hydroxylase, FLS, flavonol synthase; FNS, flavone synthase; C3H, coumarate 3-hydroxylase; COMT, caffeoyl/CoA-3-O-methyltransferase; CCR, cinnamoyl-CoA reductase; HQT, hydroxycinnamoyl-CoA quinate transferase; CAD, cinnamoyl-alcohol dehydrogenase; POD, peroxidase, HCT, hydroxycinnamoyl-Coenzyme A shikimate/quinate hydroxycinnamoyltransferase. Metabolites know to be absent in *P. patens* are in grey. Metabolic steps in the shikimate and phenylpropanoid pathways are indicated with yellow and blue arrows, respectively. Adapted from Tohge et al (2013) and Yeh et al. (2014). **(b)** Validation of differentially expressed genes by RT-qPCR at MR and LR. The expression levels in *B. cinerea*-inoculated plants are relative to the corresponding level of expression in water-treated plants at the indicated time points. Ubi was used as the reference gene. Results are reported as means ± standard deviation (SD) of three samples for each treatment. Asterisks indicate a statistically significant difference between *B. cinerea*-inoculated and the water-treated plants (Students t-test, *P < 0.01; **P < 0.001).

### *P. patens* induces the expression of genes involved in pathogen perception, signaling and transcriptional regulation during *B. cinerea* infection

Pathogen perception by membrane-localized RLKs is essential for proper activation of plant defense mechanisms. In total, 55 *RLK* genes were differentially expressed in *B. cinerea-* inoculated tissues compared to control plants, mainly Leucine rich repeat (LRR) containing proteins (including interleukin-1 receptor-associated kinases), as well as putative LysM-containing receptors, lectin receptor kinases and a proline-rich receptor-like protein kinase (**Fig. 5; Supplementary Table S9**). Upregulation of 21 *RLKs* at MR and 24 *RLKs* at LR was observed, which corresponded to for 91% and 49% of the differentially expressed RLKs at each time point, respectively. In addition, genes encoding proteins known to associate to RLKs in angiosperms, such as PTI1-like tyrosine-protein kinase, Mildew resistance Locus (MLO), and putative U-BOX E3 ubiquitin ligases involved in modulating FLS2-mediated immune signaling (García et al. 2014; Zhou and Zeng 2018), were all upregulated in response to *B. cinerea* inoculation. Moreover, expression of three putative R protein-encoding genes was affected by *B. cinerea,* including a Toll/interleukin-1 like Receptor (TIR) domain, a NBS-LRR containing proteins and a broad-spectrum mildew resistance protein.

**Fig. 5:**
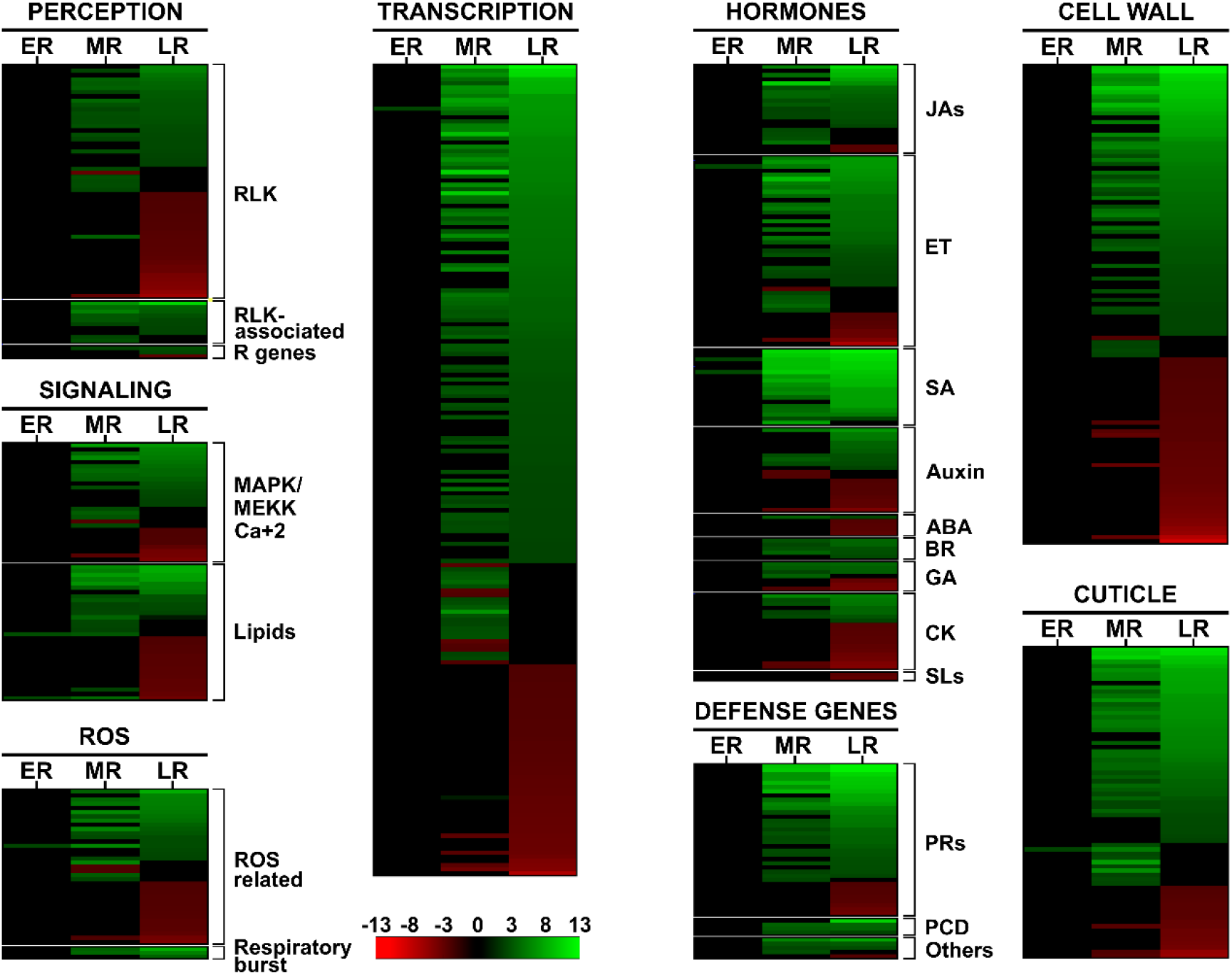
Heatmap representations of differentially expressed *P. patens* genes during *B. cinerea* infection at ER, MR and LR. *P. patens* DEGs values with log2 FC ≥2.0 or ≤ −2.0 and FDR≤ 0.05 were used for heatmap construction and grouped according to their role in defense. Normalized values relative to control and ordered according to LR log2 FC (up to downregulated), are shown in a green-red scale. Abbreviations: RLKs, receptor like kinases; R, resistance; MAPK, Mitogen-activated protein kinase; MEKK, Mitogen-activated protein kinase kinase kinase; ROS, reactive oxygen species; JAs, jasmonates; ET, ethylene; SA, salicylic acid; ABA, abscisc acid; BR, brassinosteroids; GA, gibberellic acid; CK, cytokinin; SLs, stringolactones, PR, pathogenesis-related; PCD, programmed cell death. See Supplementary Table S9-S11 for complete information.

In angiosperms, recognition of pathogens triggers Ca^2+^ fluxes, a rapid production of ROS via consumption of oxygen in a so-called oxidative burst, phosphorylation events by MPKs or Ca2+ dependent protein kinases (CDPKs), and transcriptional reprogramming (Knogge et al. 2009). In our transcriptomic analysis*, B. cinerea*-responsive DEGs related to Ca^2+^ signaling encode 24 proteins (17 upregulated), including calmodulin, calcium-binding protein (CML) and calcium dependent protein kinases. ROS are rapidly produced in moss cells after *B. cinerea* infection (Ponce de Leon et al. 2012), and here we show that transcript levels of genes involved in oxidative burst such as respiratory burst oxidase (*RBOH*) and NAD(P)H oxidase are highly induced during *B. cinerea* infection. In addition, *B. cinerea* inoculation led to differential expression of genes encoding calcium-dependent protein kinase (CDPK), MPK and mitogen-activated protein kinase kinase kinase (MEKK). Moreover, transcript levels of all 18 *P. patens* DEGs encoding enzymes involved in signaling and synthesis of PA and sphingolipids increased at MR, including lipid phosphate phosphatase, phosphatidylinositol phospholipase C, phospholipase D and A, diacylglycerol kinase and glucosylceramidase. At LR, of the 28 DEGs related to these lipid messengers, almost 50 % were upregulated, suggesting their involvement in moss defense against *B. cinerea* (**Fig. 5; Supplementary Table S9**).

TFs play a central role in the activation or suppression of gene expression during plant immunity. Consistently, TF activity was a MF GO term overrepresented in upregulated genes at MR and LR (**Supplementary Table S6**). In total, 192 *P. patens* genes encoding TFs and proteins related to transcription were differentially expressed during *B. cinerea* infection (**Fig. 5; Supplementary Table S9**). Apetala2/Ethylene Responsive Factor (AP2/ERF) was the largest family in both MR and LR. In total, 44 *AP2/ERF* were responsive to *B. cinerea;* 59% and 68% were upregulated at MR and LR, respectively. Moreover, almost all WRKYs, 17 of 19 DEGs, were upregulated during *B. cinerea* infection, as well as 9 VQ motif-containing protein encoding genes, which are transcription regulators that interact with WRKYs to modulate downstream gene expression involved in plant immunity and response to abiotic stress (Jing and Lin 2015). The other *B. cinerea*-responsive *P. patens* DEGs encoded for TFs know to regulate angiosperm responses to biotic stress comprised MYB, bHLH, B3, NAC, GRAS, and PLATZ, as well as other TFs. Taken together, these results indicate that pathogen perception, signaling and activation of TFs leading to defense mechanisms in early-diverged land plants and angiosperms are in general conserved.

### Involvement of hormonal pathways in the *P. patens* defense response against *B. cinerea*

In angiosperms, the plant hormones SA, JA, ethylene (ET), ABA, auxins, cytokinins (CK), gibberellins (GA), and brassinosteroids (BR) function in complex networks to regulate plant resistance against pathogens (Denancé et al. 2013). In this study, the involvement of jasmonates and ET during the defense response of *P. patens* to *B. cinerea* infection was highlighted by overrepresented DEGs (**Fig. 5; Supplementary Table S10**). In total, 19 of 21 DEGs related to jasmonates synthesis and signaling were upregulated during MR and LR, including lipoxygenases (LOXs), allene oxide synthase (AOS), allene oxide cyclases (AOC), OPDA reductase (OPR), OPC-8:0 CoA ligase 1 (OPCL1), TIFY (JAZ;JASMONATE ZIM DOMAIN), and TFs bHLH (bHLH27 and EGL1). This finding is consistent with previous results showing induced expression of *LOXs*, *AOS* and *OPR*, and increased OPDA levels during infection of *P. patens* tissues with *B. cinerea* (Ponce de León et al., 2012). In addition, 34 of 44 DEGs belonging to the *AP2/ERF* transcription factor family, as well as a putative ET receptor (ETR; two-component sensor histidine kinase) gene, were upregulated in response to *B. cinerea* inoculation.

SA is synthesized in plants from cinnamic acid via PAL or from chorismate via isochorismate synthase (ICS) catalyzed steps. Here, we show that expression of 17 PAL-encoding genes and a gene encoding a putative SIZ1 protein involved in SA signaling during innate immunity (Lee et al. 2007), increase during *B. cinerea* infection. Consistently, SA levels rise in *P. patens* during *B. cinerea* infection, suggesting that this hormone participate in moss defense against biotic stress (Ponce de León et al. 2012). Other DEGs related to auxin, CK, ABA, BR, stringolactones and GA biosynthesis and signaling were differentially expressed (**Fig. 5; Supplementary Table S10)**, although their role in defense against *B. cinerea* was uncertain since some genes were upregulated by the fungus, while others were downregulated. For example, genes encoding GH3 that conjugate amino acids to auxin and other hormones, and several auxin responsive genes were downregulated after *B. cinerea* infection, which is consistent with reduced levels of auxin in *B. cinerea*-infected *P. paten*s tissues (Ponce de León et al. 2012). However, auxin inducible proteins and auxin transporters encoding genes were also upregulated by *B. cinerea*. Collectively, these results indicate that like in angiosperms complex hormonal pathways operate during moss defense response against *B. cinerea*.

### *P. patens* activates expression of genes with conserved functions in plant defense against pathogens

Induced expression of PRs, which are small proteins with antimicrobial activities, is one of the typical hallmarks of defense response activation against pathogens (van Loon et al. 2006). In total, 23 and 27 of 36 DEGs encoding PR, were upregulated at MR and LR, respectively, including genes encoding PR-1 (small cysteine-rich secreted protein), PR-2 (ß-1,3 glucanases), PR-3 (chitinase), *PR-5* (thaumatin), PR-9 (peroxidases), and PR-10 (Bet v I), (**Fig. 5; Supplementary Table S11**). Peroxidases were among the family with most *B. cinerea*-inducible gene members, including Prx34 (Pp3c4_14530), which participates in *P. patens* defense against pathogens (Lehtonen et al. 2009). Other *B*. cinerea-responsive DEGs related to oxidative stress, encoded glutathione S-transferase (GST), peroxiredoxin, thioredoxin, ferredoxins, catalase and superoxide dismutase. Three TSPO genes were upregulated in *B. cinerea*-infected *P. patens* tissues, including Pp3c24_14170, which is involved in redox homeostasis in moss cells (Lehtonen et al. 2012). Similarly, a gene encoding a tocopherol cyclase, which is involved in production of tocopherol in plastids where it protects membranes from oxidative degradation by ROS (Stahl et al., 2019), and a chloroquine-resistance transporter-like involved in glutathione homeostasis during defense against pathogens (Maughan et al. 2010), were also upregulated during *B. cinerea* infection. ROS accumulation can lead to an HR response, which is a type of programmed cell death (PCD) (Lamb and Dixon 1997). We have previously observed that an HR-like response is triggered in *B. cinerea* infected *P. patens* tissues, which probably facilitates colonization of moss tissues (Ponce de León et al. 2012). Here we show that several genes involved in PCD were upregulated in *P. patens* during *B. cinerea* colonization, including genes encoding subtilisin-like proteases, metacaspase, and a E3 ubiquitin-protein ligase BOI that was previously shown to suppress pathogen-induced cell death in angiosperms (Luo et al. 2010) (**Fig. 5; Supplementary Table S11).**

Plant cuticle and cell walls serve as the first line of defense against *B. cinerea* limiting the advancement of growing hyphae in angiosperms (AbuQamar et al. 2016). We observed by microscopic analysis that at MR plant cells start to modify their cell walls in response to *B. cinerea* colonization (**Fig. 1**). Consistently, 113 *B. cinerea*-responsive DEGs were associated to cell wall biosynthesis or modification. Several genes encoding enzymes involved in cellulose, hemicellulose, arabinose and pectin biosynthesis were induced after *B. cinerea* infection, including cellulose synthase, glucuronosyl transferase, UDP-glucuronate 4-epimerase, UDP-glucuronate decarboxylase and UDP-arabinose 4-epimerase. Other DEGs encode enzymes that participate in endogenous cell wall degradation such as endo-glucanases, endo-xylanase, pectin esterases and pectate lyases, and cell wall modifications such as pectin methylesterase inhibitor and expansins (**Fig. 5; Supplementary Table S11**). In addition, a high number of genes involved in cuticle lipid biosynthetic pathways (Lee et al., 2020), were differentially regulated in response to this pathogen. Most genes were upregulated, including several genes encoding ATP citrate synthases, acetyl-CoA carboxylases, Long Chain Acyl CoA synthetase, Ketoacyl CoA synthases, and other enzymes involved in very-long-chain fatty acids (VLCFA) and cutin synthesis (**Fig. 5; Supplementary Table S11**). Moreover, other *B. cinerea*-responsive DEGs encode proteins with known functions in angiosperm defense against pathogens, including proteins that recognize or modify chitin from the pathogen, an α-dioxygenase (α-DOX), and different types of transporters (**Supplementary Table S3**). Thus, our results suggest that conserved defense mechanisms were likely present in the common ancestor of land plants.

### Genes acquired by horizontal transfer and orphan genes participate in moss defense against *B. cinerea*

*P. patens* has nuclear genes acquired by horizontal gene transfer from bacteria, viruses and fungi (Yue et al. 2012). The acquired functions probably played a role in the transition of plants from aquatic to terrestrial environments as well as defense mechanisms against pathogens. Of the 128 horizontal transferred genes identified in Yue et al. (2012), 18 were present in our RNAseq analysis, including DEGs encoding acyl-activating enzyme, glutamate-cysteine ligase, acid phosphatase, allantoate deiminase, methionine-gamma-lyases, oleoyl-[acyl-carrier-protein] hydrolase and beta-glucosidases (**Supplementary Table S12**). Several of these genes could contribute to defense against *B. cinerea* since they are involved in defense processes such as synthesis of auxin, glutathione, as well as degradation of purine, cellulose and methionine. While these genes were probably acquired from bacteria and have homologs in *Arabidopsis*, other DEGs such as genes encoding a UBIA prenyltransferase and phosphoglycerate kinase, involved in menaquinone biosynthesis and glycolysis respectively, were probably obtained from δ-proteobacteria and do not have homologs in *Arabidopsis*. A gene encoding a MFS transporter involved in sugar transport and two heterokaryon incompatibility proteins (HET), which are only present in fungi and moss, were upregulated during *B. cinerea* infection. Interestingly, a sea anemone cytotoxic protein-coding gene acquired probably by cnidarian and involved in dehydration stress responses of *P. patens* (Hoang et al. 2009), was also upregulated with *B. cinerea*. This gene is JA, SA, ET and ABA inducible, and although the exact function is at present unknown, it could have cytotoxic properties against pathogens.

Since almost 50% of the *B. cinerea*-responsive genes obtained by horizontal transfer have no homologues in *Arabidopsis* according to Yue et al. (2012), we looked in more detail at DEGs with no homologues in other plant species. A high number of *B. cinerea*-responsive DEGs, 599 genes representing 19 % of the total DEGs, encode putative proteins that do not have significant sequence similarity (e-value >1.0E^-3^) to any other Viridiplantae proteins/peptides and were considered orphan genes (Wilson et al. 2005). Among the total orphan genes, three, 198 and 236 were upregulated at ER, MR and LR, respectively, while 50 were downregulated at MR and 274 at LR (**Fig. 6; Supplementary Table S12**). Most of these *B. cinerea-* responsive species-specific genes (92%) are described as with unknown functions in the Phytozome database. Of the 599 orphan genes, 376 showed no blast hits, meaning that they do not have any homologs and were *de novo* created genes. In addition, 137 orphan genes belong to multimember gene families, and 12 were not expressed in the *P. patens* Atlas project (Perroud et al. 2018) (**Fig. 6**). Two *B. cinerea* inducible orphan genes were secreted from moss tissues after chitosan treatment (Lehtonen et al. 2014), including Pp3c19_19780, Pp3c18_6260, suggesting that they play a role in moss defense. Taken together, the acquisition of horizontal transferred gene as well as orphan genes could have provided adaptive advantages to encounter microbial colonization in basal land plants.

**Fig. 6:**
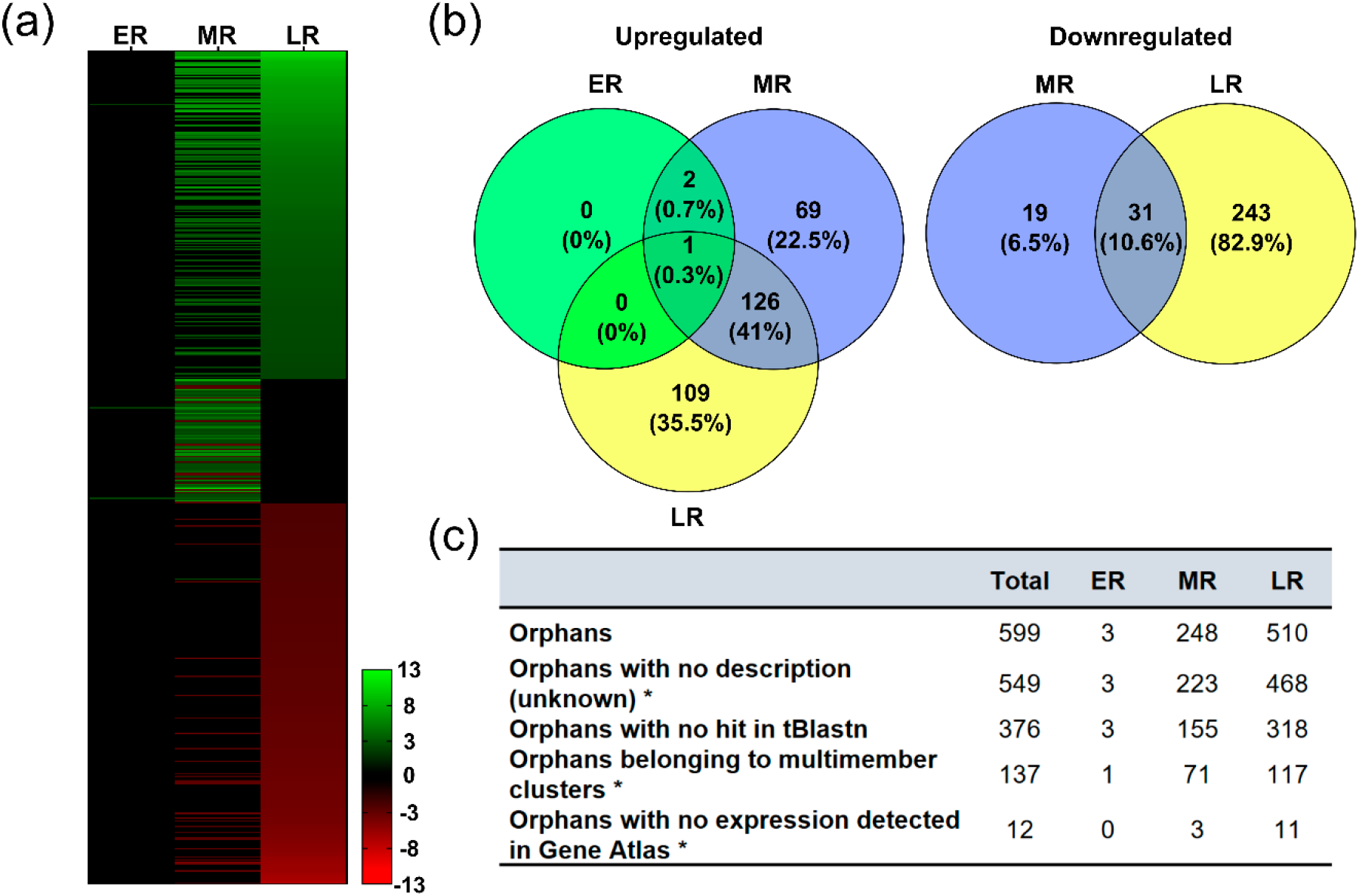
*P. patens* orphan genes expression in response to *B. cinerea*. **(a)** Heatmap showing the distribution of *P. patens* orphan DEGs with log2 FC ≥2.0 or ≤ −2.0 and FDR≤ 0.05 during *B. cinerea* infection at ER, MR and LR. Normalized values relative to control and ordered according to LR log2 FC (up to downregulated), are shown in a green-red scale. **(b)** Venn diagrams showing overlap of expressed orphan genes (upregulated and downregulated) at ER, MR and LR. **(C)** Distribution of identified *B. cinerea*-responsive *P. patens* orphan genes and some orphan sequence characteristics. Asterisks indicate information extracted from phytozome.org. See Supplementary Table S12 for complete information.

## Discussion

With the conquest of land, plants developed adaptation mechanisms to cope with new pathogens present in air and soil. The identification of the genetic basis of such innovations is relevant for understanding how plant defense responses developed during the interaction with microbial pathogens across the green lineage. In the present study, we show that a descendant of the earliest land plants, the moss *P. patens*, responds to the fungus *B. cinerea* via a complex and integrated set of defenses encompassing both induced responses and downregulation of specific pathways. While a small group of plant genes, consisting of 17 upregulated genes, showed differential expression at ER, during MR 1.032 genes were induced by *B. cinerea,* representing 67 % of the total upregulated DEGs (1.548) present over the entire time course of infection. This finding indicates the occurrence of massive gene expression in moss tissues that precede cell invasion by fungal hyphae and the onset of disease symptoms. *P. patens* transcriptional reprogramming at MR included genes encoding proteins involved in pathogen perception, signaling, transcription, hormonal signaling, metabolic pathways and genes with diverse role in defense against biotic stress. Upregulation of more than 67% (693 genes) of the MR inducible DEGs continued at LR, where *B. cinerea* hyphae colonized the plant tissues and symptom development was visible. Interestingly, while only 150 DEGs were downregulated during MR, expression levels of 1.482 genes decreased at LR, including those related to photosynthesis, cell cycle and cell division. Downregulation of photosynthesis and associated processes in response to *B. cinerea* infection and other plant pathogens is a common phenomenon in angiosperms (Bilgin et al. 2010; Windram et al. 2012). *P. patens* could reallocate nitrogen resources for synthesis of new defense proteins as has been observed in angiosperms (Windram et al. 2012). Moreover, *B. cinerea* infection produced browning of chloroplasts, followed by the breakdown of these organelles (Ponce de León et al. 2007). Besides, plants balance cell cycle regulation and immune response to pathogens through manipulating the level of cyclins and cyclin-dependent kinase (Qi and Zhang 2020), which is consistent with downregulation of genes encoding these type of proteins in our transcriptomic data during *P. patens*-*B. cinerea* interaction. This finding together with the fact that chitin and crude extracts of pathogenic *Pseudomonas* and *Plectosphaerella cucumerina* treatments inhibited *P. patens* and *M. polymporha* growth, respectively (Galloto et al. 2020; Gimenez-Ibanez et al. 2019), indicate that like in angiosperms (Ranf et al. 2011), PAMPs also trigger growth inhibition in bryophytes.

Our results show that *P. patens* perceives *B. cinerea* and activates the expression of multiple defense genes associated to PTI and ETI in angiosperms, suggesting a conserved role of the encoded proteins in defense throughout the green plant lineage. Thirty upregulated DEGs encoded RLKs, mainly proteins with LRR domains, demonstrating the importance to express many receptors that may detect molecular components of the pathogen in order to mount an effective moss defense response. Consistently, RLK families expanded with the establishment of land plants as an adaptation mechanism to fast-evolving pathogens (Lehti-Shiu et al. 2009). Interestingly, while *P. patens* has a CERK1 receptor that perceives fungal chitin and bacterial peptidyl glycan leading to plant immunity (Bressendorff et al. 2016), flagellin-derived peptide flg22 and the elongation factor Tu are not sensed by moss cells since *P. patens* lacks FLS2 and EFR orthologues receptors (Boller and Felix 2009). Similarly, *M. polymorpha* does not have an FLS2 orthologue and does not respond to flg22 (Gimenez-Ibanez et al. 2019). Genes encoding proteins known to associate with RLKs in angiosperms were also upregulated in *P. patens,* including PTI tyrosine-protein kinase, MLO and U-BOX E3 ubiquitin ligases, suggesting that the common ancestor of land plants likely possessed a perception mechanism of PAMPs through different types of RLKs and associated proteins that modulate immune responses to pathogens. In plants, the second layer of immune system relies on R proteins that act inside the cells and confer recognition of pathogen effectors to trigger ETI (Jones and Dangl 2006). Although mosses have a high number of R genes that could have provided early land plants with a genetic toolkit needed for survival in a terrestrial environment (Gao et al. 2018)*, B. cinerea* infection only altered the expression of three R genes in *P. patens*. This result was unexpected since many R genes were differentially regulated in other plant-*B. cinerea* interactions (Windram et al. 2012; Haile et al. 2019). Although the activation of ETI was demonstrated during *P. patens-B. cinerea* interaction through the presence of an HR-like response (Ponce de León et al. 2012), and our transcriptomic analysis show that several genes involved in plant PCD are induced, further research is needed to understand the role played by R genes during infection of moss cells with pathogens and subsequent activation of plant immunity.

Alterations in the *P. patens* transcriptome due to *B. cinerea* colonization comprised gene members related to signaling pathways in angiosperms, including lipid messengers such as inositol 1,4,5-trisphosphate (IP3), PA, sphingolipids, and calcium. Consistently, chitin triggers cytosolic calcium increase in *P. patens* cells that leads to induced expression of defense genes (Galloto et al. 2020). Moreover, ROS level rises quickly in *P. patens* cells during *B. cinerea* infection (Ponce de León et al. 2012), acting probably as second messengers during moss immunity. Upregulation of *B. cinerea*-responsive genes encoding peroxidases, including Prx34 known to be responsible for the oxidative burst in *P. patens* after chitin treatment (Lehtonen et al. 2012), as well as a RBOH and NADPH oxidases, highlighted the role played by ROS in moss defense. In fact, mutants lacking the secreted Prx34 are more susceptible to fungal pathogens (Lehtonen et al. 2009). Our data showing increased expression of several *TSPO* genes involved in mitochondrial tetrapyrrole transport and known to control ROS levels in *P. patens* ((Lehtonen et al. 2012), as well as other oxidative stress related genes encoding peroxiredoxins, thioredoxins, ferredoxins and GSTs during *B. cinerea* infection, support the importance to maintain a redox balance through generation and elimination of ROS. This result is in accordance with increased ROS production upon pathogen recognition in *tpso1* mutants that may result in severe damage, cell death and disease symptoms after fungal infection (Lehtonen et al. 2012). Genes encoding MEKKs and MPK were differentially expressed during *B. cinerea* infection, indicating that they are involved in immune responses and may promote defense gene expression. *P. patens* has components of an immune pathway that include MEKK, MAP kinase kinase (MKK), and two MPKs that are required for defense responses to fungal chitin (Bressendorff et al. 2016). MPK moss mutants result in loss of basal resistance to pathogens, evidenced by reduced defense gene expression and cell wall associated defenses (Bressendorff et al. 2016). TFs with known functions in angiosperm defenses against *B. cinerea*, including AP2/ERF, WRKY, MYB, bHLH, PLATZ and NAC (Mengiste et al. 2003; Berrocal-Lobo et al. 2002; Zhao et al. 2012; AbuQamar et al. 2017), were also differentially expressed in fungal-infected moss tissues. Compared with algae, the numbers of members of TFs families including MYB, WRKY, and AP2/ERF have expanded significantly in land plants and might be associated to terrestrial adaptation (Rensing et al. 2008; Rinerson et al. 2015; Pu et al. 2020). The high number of AP2/ERF TFs, as well as WRKYs and associated regulatory proteins with VQ motif, upregulated during *B. cinerea* infection, suggest their involvement in defenses to counteract pathogens among these adaptation mechanisms. Previous work also showed induced expression of AP2/ERF family members by other biotic stresses such as elicitors of *P. carotovorum,* chitosan and *Phytophthora capsici* (Alvarez et al. 2016, Bressendorff et al. 2016; Overdijk et al. 2016). This result is consistent with a central regulatory role played by the AP2/ERF family during different stress conditions in *P. patens* (Hiss et al. 2014). Interestingly, three *B. cinerea*-inducible NAC TFs (Pp3c13_20650, Pp3c3_12890 and Pp3c4_20430), that are involved in *P. patens* water-conducting cell development, showed cell wall thickening and PCD when overexpressed in *P. patens* (Xu et al. 2014). Collectively, these findings indicate that defense signaling pathways were already present in early diverging plants and that TFs have an ancestral role in orchestrating a defense response during pathogen infection.

Like in angiosperms (AbuQamar et al. 2017), our findings suggest that jasmonates, ET and SA play a role in *P. patens* defense responses against *B. cinerea*. Bryophytes have conserved sequences for all jasmonoyl-isoleucine (JA-Ile) signaling pathway components but lack the hormone JA-Ile (Monte et al. 2018). Instead, dinor-OPDA is the ligand for the functionally conserved JA-Ile receptor (COI1) in bryophytes (Monte et al. 2018). In the present study, we showed upregulation of several genes encoding enzymes involved in jasmonate production, as well as JAZs and bHLH TFs upon *B. cinerea* inoculation. Moreover, in *B. cinerea*-infected *P. patens* tissues the levels of OPDA increases leading to expression of the defense gene *PAL* (Ponce de León et al. 2012), demonstrating that in spite of the difference in the identity of the bioactive hormone, their role in defense against pathogens is conserved among angiosperms and bryophytes. ET is synthesized in *P. patens*, although the key enzyme 1-aminocyclopropane-1-carboxylate (ACC) synthase (ACS) encoded by two genes, have no ACS activity (Sun et al. 2016). This finding together with evidence showing that *P. patens* and *M. polymorpha* are unable to convert exogenously added ACC to ET, indicate that a different ethylene biosynthetic pathway may exist in bryophytes (Sun et al. 2016). ET is involved in *P. patens* adaptive responses to drought and submergence through the action of ET receptor (ETR) (Yasumura et al. 2012). In our analysis, transcript levels of an ETR and several AP2/ERFs encoding genes increased during *B. cinerea* infection, indicating that as in angiosperms, ethylene is probably involved in defense responses against biotic stress. However, further research is required to determine the role of this hormone during pathogen infection of early-diverging land plants.

SA increases in *P. patens* tissues after *B. cinerea* inoculation leading to induced expression of *PAL,* indicating that SA is synthesized and perceived in moss cells (Ponce de León et al. 2012). Similarly, SA levels increase in *M. polymorpha* after *Pseudomonas* extracts treatment, leading to activation of typical SA markers such as *PR-1, PR-2,* and *PR-5* (Gimenez-Ibanez et al. 2019). Thus, bryophytes perceive microbial pathogens and activate SA signaling to restrict pathogen colonization similarly as in angiosperms. *P. patens* has only one SA receptor, NPR1-like protein, which partially rescue *Arabidopsis npr1* mutant, suggesting the presence of a NPR1-like positive regulator of SA-mediated defense responses in mosses (Peng et al. 2017). The SA pathway was probably present in the common ancestor of land plants since all genes of this pathway are present in bryophytes but not in algae (Wang et al. 2015). However, the recent finding that *P. patens* NPR1-like receptor did not complement *Arabidopsis npr3-4* double mutant suggests that fine-tuning of SA-induced defense responses probably evolved later in angiosperms (Peng et al. 2017).

Other DEGs related to auxin, CK, ABA, and BR signaling show that a complex hormonal network regulate *P. patens* defense against *B. cinerea*. Auxin levels decreased in *P. patens* after *B. cinerea* infection, while ABA levels increases at 24 hpi, although this hormone could be of fungal origin used as a mechanism to contribute to infection (Ponce de León et al. 2012). BR are important for rose defense against *B. cinerea* (Liu et al., 2018), and here we show that all fungal-responsive DEGs of this pathway are upregulated in *P. patens*. However, the BR receptor and the key regulator BKI1 (Brassinosteroid Insensitive1 Kinase Inhibitor) are not present in *P. patens* and moss cells do not respond to BR (Prigge et al. 2010; Wang et al. 2015). In addition, *P. patens* lacks a GA biosynthetic pathway downstream of ent-kaurenoic acid (KA), indicating that KA metabolites instead of GA may play physiological roles (Miyazaki et al. 2018). The observation that genes encoding ent-kaurene synthases and a KA 2-oxidase (PpKA2ox; Pp3c17_7090; Miyazaki et al. 2018) were upregulated during *B. cinerea* infection, suggests a possible involvement of KA metabolites in plant defense to pathogen invasion. Collectively, these results show that ancient hormonal signaling, several of which have different features with angiosperms, are involved in *P. patens* defense responses against pathogens.

The shikimate and phenylpropanoid pathways played a crucial role in adaptation mechanisms that have evolved in early land plants to cope with abiotic stresses such as UV radiation and desiccation, as well as microbial attack (Emiliani et al. 2009). Our data highlight the importance of these pathways in moss defense against pathogens since a very high number of biosynthetic and regulatory genes were upregulated during *B. cinerea* infection. Similar results were obtained during angiosperms-*B. cinerea* interactions (Windram et al. 2012; Haile et al. 2019, 2020). The contribution of phenylpropanoids to bryophyte defense against pathogens was evidenced in several other *P. patens*-pathogen interactions (Oliver et al. 2009; Reboledo et al. 2015: Alvarez et al. 2016), and during infection of the liverwort *M. polymorpha* with the oomycete *P. palmivora* (Carella et al. 2019). *P. paten*s genome duplicated 45 Mya and several gene families expanded, including *PALs* and *CHS*s (Koduri et al. 2010; Wolf et al. 2010). The presence of different CHSs activities in *P. patens* (Li et al., 2018), and the fact that most *PALs* and *CHSs* are induced during *B. cinerea* infection, suggest that plasticity of secondary metabolism could have allowed early land plants to adapt more easily to pathogens. In angiosperms, the phenylpropanoid pathway contributes with a vast array of metabolites with different roles in defense against invading pathogens, such as preformed chemical and physical barriers (cuticle and lignin), antimicrobial activities (lignans, coumarins, isoflavonoids and flavonoids), as well as signaling molecules (like SA) involved in expression of defense genes in local and systemic tissues (Dixon et al. 2002). Cinnamic acid levels increases after *P. carotovorum* elicitor treatment (Alvarez et al. 2016), and caffeic acid, coumaric acid, 4-hydroxybenzoic acid and caffeoyl quinic acid have been detected in *P. patens* tissues, some of which have antimicrobial properties (Erxleben et al. 2012; Richter et al. 2012). Several *B. cinerea*-responsive DEGs were annotated as CHI, F3H and FLS. However, unlike angiosperms, *P. patens* does not seem to contain enzymes required for the synthesis of flavonones and flavones. Both CHIs are enhancers of flavonoid production genes encoding type IV CHI proteins with no CHI activity (Ngaki et al. 2012; Waki et al. 2020), and there are no evident orthologs for F3H and FLS (Koduri et al. 2010; Kawai et al. 2014). Thus, *P. patens* probably lacks the later steps in flavonoid biosynthesis, in contrast to *M. polymorpha*, which accumulates different types of flavonoids (Albert et al. 2018). *P. patens* has a pre-lignin pathway instead of true lignin (Weng and Chapple, 2010), and we showed previously that polyphenolic compounds reinforce the cell wall upon *B. cinerea* infection (Ponce de León et al. 2012). Upregulation of DIR, PPO and Prx encoding genes, which are involved in monolignols coupling, may contribute to strengthen the polyphenolic cell walls. This finding together with the fact that SA levels rises upon *B. cinerea* inoculation (Ponce de León et al. 2012), suggests that cinnamic acid-derived metabolites and monolignol-containing compounds are important for moss defense against pathogens. Recently, Renault et al. (2017) showed that the pre-lignin pathway of *P. patens* involves the formation of caffeic acid coupled to the hexose-derived ascorbate pathway, rather than the shikimate pathway, as is reported for angiosperms. This leads to the formation of soluble caffeoyl-threonic acids, which are essential for the synthesis of the moss cuticle. In addition, transcript levels of several *B. cinerea*-responsive MYB, belonging to R2R3 MYB (Pu et al., 2020), and bHLB TFs are involved in phenylpropanoid biosynthesis in angiosperms (Xu et al. 2013). Consistently, a functional analysis demonstrated that a R2R3-MYB TF activates flavonoids biosynthesis in *M. polymorpha*, and the expression of members of this pathway is necessary for liverwort defense against *P. palmivora* (Carella et al. 2019). Collectively, these observations indicate that like in angiosperms, phenylpropanoids play important roles in bryophyte defense, suggesting that the emergence of specialized secondary metabolites in early land plants was probably important to protect plant tissues from invading microorganisms.

PRs belong to a protein family with demonstrated roles in defense responses of angiosperms to pathogens (van Loon et al. 2006). Homologs of most PR gene members are present in bryophytes with the exception of *PR-12* (defensin) and *PR-13* (thionin) that probably evolved later in angiosperms (Visser et al. 2018). In our transcriptomic analysis, *PR-1*, *PR-2*, *PR-3*, *PR-5, PR-9* and *PR-10* were upregulated during *B. cinerea* infection, indicating their contribution to moss defense. Consistently, constitutive expression of PR-10 (Pp3c2_27350) increased resistance to the oomycete *P. irregulare* in *P. patens*, where cell wall reinforcement probably contributes to halt pathogen colonization (Castro et al. 2016). Overexpression of *P. patens* PR-10 also increased resistance to this oomycete in the evolutionary distant plant *Arabidopsis* (Castro et al. 2016). Peroxidases encoding DEGs were among the highest *B. cinerea*-inducible members of the moss PR family, similarly as in *P. palmivora*-infected *M. polymorpha* tissues (Carella et al. 2018, 2019). *P. patens* mutants affected in *Prx34* displayed an enhanced susceptibility to pathogenic and saprophytic fungi (Lehtonen et al. 2009), highlighting the involvement of peroxidases in moss defense. In addition, *B. cinerea*-inducible Prx34, *PR-5* and a thioredoxin (Pp3c6_10090), were secreted to the apoplast in chitosan-treated moss tissues (Lehtonen et al. 2014). These apoplastic proteins could contribute to moss defense by regulating cellular redox status or exerting antifungal activities, and/or through cell wall reinforcement by catalyzing H2O2-dependent cross-linking of cell wall components. The importance of cell wall defenses in *P. patens* response to *B. cinerea* colonization was evidenced by differential expression of a high number of genes involved in cell wall synthesis and modifications. Two genes encoding pectin methylesterase inhibitor were upregulated at MR and LR, which are involve in making the wall more resistant to degradation by fungal enzymes by increasing methylated pectin in the cell wall (Lionetti et al. 2007). Moreover, several downregulated genes encoded expansins, known to influence cell wall extensibility and *B. cinerea* susceptibility in *Arabidopsis* (AbuQamar 2013). In addition, genes involved in cuticle biosynthetic pathways, including those related to fatty acid, VLCFA, cutin and wax synthesis (Lee at al. 2020), were mostly upregulated during *B. cinerea* infection. Cell wall reinforcement and cuticle associated defenses probably serve as the first line of defense to limit the advancement of growing hyphae, like in angiosperms (AbuQamar et al. 2016). Several LOXs and an α-Dioxygenase encoding genes were upregulated upon *B. cinerea* inoculation, and similarly as in angiosperms they produce oxylipins with different roles in plant defense against pathogens (Ponce de León et al. 2015). α-DOX-derived oxylipins increases after pathogen infection in *P. patens* and *Arabidopsis* and protect plant tissues against cellular damage caused by pathogens (Ponce de León et al. 2002; Machado et al. 2015). Collectively, these findings suggest a conserved role for a high number of DEGs in mediating defense responses to fungal pathogens throughout the green plant lineage.

Several genes present in our *P. patens-B. cinerea* transcriptomes and associated to defense were acquired by horizontal gene transfer from prokaryotes and fungi. Upregulated DEGs included genes involves in auxin, glutathione, fatty acid biosynthesis, glycolysis and sugar transport (Yue et al. 2012), which participate in plant disease resistance. Moreover, a UBIA prenyltransferase encoding gene from proteobacterial origin was upregulated by *B. cinerea* infection, which could be involved in ubiquinone synthesis that has antioxidant activity (Li 2016). Interestingly, two *B. cinerea* inducible genes encoded heterokaryon incompatibility proteins (HET) obtained probably from fungi, which participate in innate immune response by protecting the fungus from pathogens and serve as damaged self-recognition system by inducing PCD (Medina-Castellanos et al 2018). Fungal colonization played a fundamental role in land colonization by plants (Remy et al., 1994), and gene acquisition following fungal interaction might have represented adaptive benefits for resistance to pathogen colonization. In addition, *P. patens* has 13 % orphan genes, which are species-specific genes with no orthologs in other plants (Zimmer et al. 2013). Our findings show that a large proportion of *B. cinerea*-responsive *P. patens* DEGs (599 genes: 19%) correspond to orphan genes, from which 376 were *de novo* created genes, suggesting that they could represent innovative adaptive strategies to biotic stress. The involvement of orphan genes in cold acclimation (Beike et al. 2015), indicate that they also participate in abiotic stress tolerance. Moreover, twelve *B. cinerea*-responsive *P. patens* orphan genes were not expressed in any condition present in the large-scale RNA-seq *P. patens* Altlas project (Perroud et al. 2018), suggesting that they could represent pathogen-specific orphans. Deciphering the roles played by orphan genes, most of which have no annotations, during biotic and abiotic stress in *P. patens*, will help to understand how they contributed to stress adaptation of early-diverging land plants. In conclusion, our results reveal that conserved defense mechanisms between extant bryophytes and angiosperms, as well as moss-specific defenses are part of the *P. patens* immune system, which could have been pivotal for land colonization by plants. Further studies on *P. patens*-pathogen interactions will contribute to uncover the molecular mechanisms underlying moss-specific defenses and their involvement during coevolution of ancient land plants and pathogens.

## Availability of supporting data

The sequencing raw data from the RNA-Seq libraries were deposited on the Sequence Read Archive from NCBI under SRA accession: PRJNA647932. Data are available through https://www.ncbi.nlm.nih.gov/bioproject/647932. In addition, data sets supporting the results of this article are included in Additional files.

## Acknowledgements

This work was supported by “Fondo Conjunto” Uruguay-México (AUCI-AMEXID), “Agencia Nacional de Investigación e Innovación (ANII) (graduate fellowships)” Uruguay, “Programa de Desarrollo de las Ciencias Básicas (PEDECIBA)” Uruguay, and “Programa para Grupo de I+D Comisión Sectorial de Investigación Científica, Universidad de la República”, Uruguay. The authors thank Héctor Romero and Andrea Zimmers for advice.

## Author contributions

GR performed the experiments; GR, AA and IPDL analyzed and interpreted the data and participated in discussions; GR and AA helped to write the article; LV participated in the interpretation of the data, artwork, discussions and drafting of the work; RM participated in the drafting of this work; all the authors revised and contributed to the final version of the manuscript; IPDL wrote the manuscript. All authors read and approved the final version of the manuscript.

## Supplementary Tables

**Supplementary Table S1: List of qPCR primers used in this study.**

**Supplementary Table S2. Summary of mapped reads of the RNA-Seq libraries.** C4, water treated plants at 4 hpi; I4, *B. cinerea* inoculated plants at 4 hpi; C8 water treated plants at 8 hpi; I8, *B. cinerea* inoculated plants at 8 hpi; C24, water treated plants at 24 hpi; I24, *B. cinerea* inoculated plants at 24 hpi; 1-3 indicate the three biological replicates at the indicated time points.

**Supplementary Table S3: List of *P. patens* differentially expressed genes (DEGs) during *B. cinerea* infection at Early Response (ER), Middle Response (MR) and Late Response (LR).**

**Supplementary Table S4: List of *P. patens* DEGs at Early Response (ER).** Expression levels of these genes are also shown at Middle Response (MR) and Late Response (LR).

**Supplementary Table S5: Enriched gene ontology (GO) terms (over-representation) for biological processes and molecular function at Middle Response (MR) and Late Response (LR).**

**Supplementary Table S6: Hierarchical clustering using REVIGO to remove functional and semantic redundancies within the significantly enriched GO terms associated with the differentially expressed genes during *B. cinerea* infection.** Significance of the over-representation of the GO term (-log10 FDR) is shown.

**Supplementary Table S7: List of differentially expressed *P. patens* genes encoding for proteins involved in the shikimate and phenylpropanoid pathways during *B. cinerea* infection at Early Response (ER), Middle Response (MR) and Late Response (LR).**

**Supplementary Table S8. List of genes used to validate the RNA-Seq expression values by qRT-PCR assay. Gene identification, fold change from RNA-seq and qRT-PCR gene and description are provided.**

**Supplementary Table S9. List of differentially expressed *P. patens* genes encoding for proteins involved in perception, signaling and transcription at Early Response (ER), Middle Response (MR) and Late Response (LR).**

**Supplementary Table S10: List of hormone related differentially expressed genes in *P.patens* at Early Response (ER), Middle Response (MR) and Late Response (LR).**

**Supplementary Table S11: List of Pathogenesis-related (PR) and other defense genes differentially expressed in *P. patens* at Early Response (ER), Middle Response (MR) and Late Response (LR).**

**Supplementary Table S12: List of differentially expressed orphan genes in *P. patens* at Early Response (ER), Middle Response (MR) and Late Response (LR).**

